# Impaired social contacts with familiar anesthetized conspecific in CA3-restricted BDNF knockout mice

**DOI:** 10.1101/168229

**Authors:** Wataru Ito, Howard Huang, Vanessa Brayman, Alexei Morozov

## Abstract

Familiarity is the vital characteristic conveyed by social cues to determine behaviors towards conspecific. Here we characterize social contacts to familiar vs unfamiliar male conspecific, anesthetized to eliminate inter-male aggression. During initial 10 min (phase-1), subjects contacted demonstrators vigorously regardless of familiarity. During subsequent 80 min (phase-2), however, they contacted more with familiar than unfamiliar conspecifics. Then, this test was applied on highly aggressive mice with hippocampal CA3-restricted BDNF knockout (KO), in which aggression may mask other behaviors. KO showed less preference to contacting familiar conspecific than wild type (WT) during phase-2 but no differences during phase-1. Among non-social behaviors, eating duration was shorter in the presence of familiar than unfamiliar conspecific in WT, but same in KO. Additionally, KO exhibited reduced pain sensitization. Altogether, these findings suggest that KO has deficits in circuits that process social cues from familiar conspecifics and pain and, possibly, underlie empathy-like behaviors.

## Introduction

When conspecifics encounter each other, the social cues that inform about familiarity are likely to trigger adaptive behaviors, which are crucial for their survival. To investigate brain circuits that process social cues, a behavioral paradigm is necessary that is sensitive enough to compare social interactions between familiar and unfamiliar conspecifics without a disruption by competing behaviors. Several tests have been established in rodents for measuring distinct social traits, including sociability (Moy et al., 2004), social memory (Ferguson et al., 2000), social transmission of food preference (Galef and Wigmore, 1983), aggression (Winslow and Miczek, 1983), dominance (Sa-Rocha et al., 2006) and empathy-like behaviors (Ben-Ami Bartal et al., 2011; Chen et al., 2009; Langford et al., 2006; Panksepp and Lahvis, 2011). Since in these tests, the subjects encounter active conspecifics, the behaviors are the function of reciprocal interactions, during which the subject and demonstrator influence one another. This bi-directionality increases the variability of behavioral readout and possibly masks certain behavioral traits. As an extreme case, the high inter-male aggression in rodents overshadows other forms of social interactions between unfamiliar males.

To overcome such limitations, we characterize the interaction with an anesthetized conspecific that eliminated both the reciprocal exchange of social cues and inter-male aggression. The anesthetized demonstrator remains a source of strong social signals, which have been found to elicit defensive responses including ultrasound vocalizations in rats (Blanchard et al., 1986; Blanchard et al., 1993).

In this study, we examine mice with the CA3-restricted knockout of BDNF, which exhibit elevated aggression and dominance towards cage mates but normal cognition and social memory (Ito et al., 2011). As predicted, the new test allowed comparisons between responses to social cues from familiar versus unfamiliar conspecific while avoiding aggression. To this end, we find a distinct social trait - sustained contacting the familiar, but not unfamiliar anesthetized conspecific - and that trait was compromised in the BDNF KO mice, which showed normal sociability in the three chamber test (Moy et al., 2004).

## Results

### Effect of BDNF CA3 KO on contacting familiar and unfamiliar demonstrator

The KO and WT mice were presented with an anesthetized demonstrator, either the sibling cage mate (familiar) or a stranger on the 129SvEv background (unfamiliar) (Fig.1A). The demonstrator was placed at the center of the cage and the cotton nest was at the corner. Subjects did not exhibit aggression, neither did they huddle; however, in the case of familiar demonstrators, they started huddling once the anesthesia wore off and demonstrator began to move, typically, after 90 min of immobility. We first analyzed physical contacts towards the anesthetized demonstrator. The “contacting” included sniffing head and genitals, allogrooming, head-to-head contact, touching any body part, sticking a nose under the body, and digging wood chip bedding underneath.

**Fig. 1.**
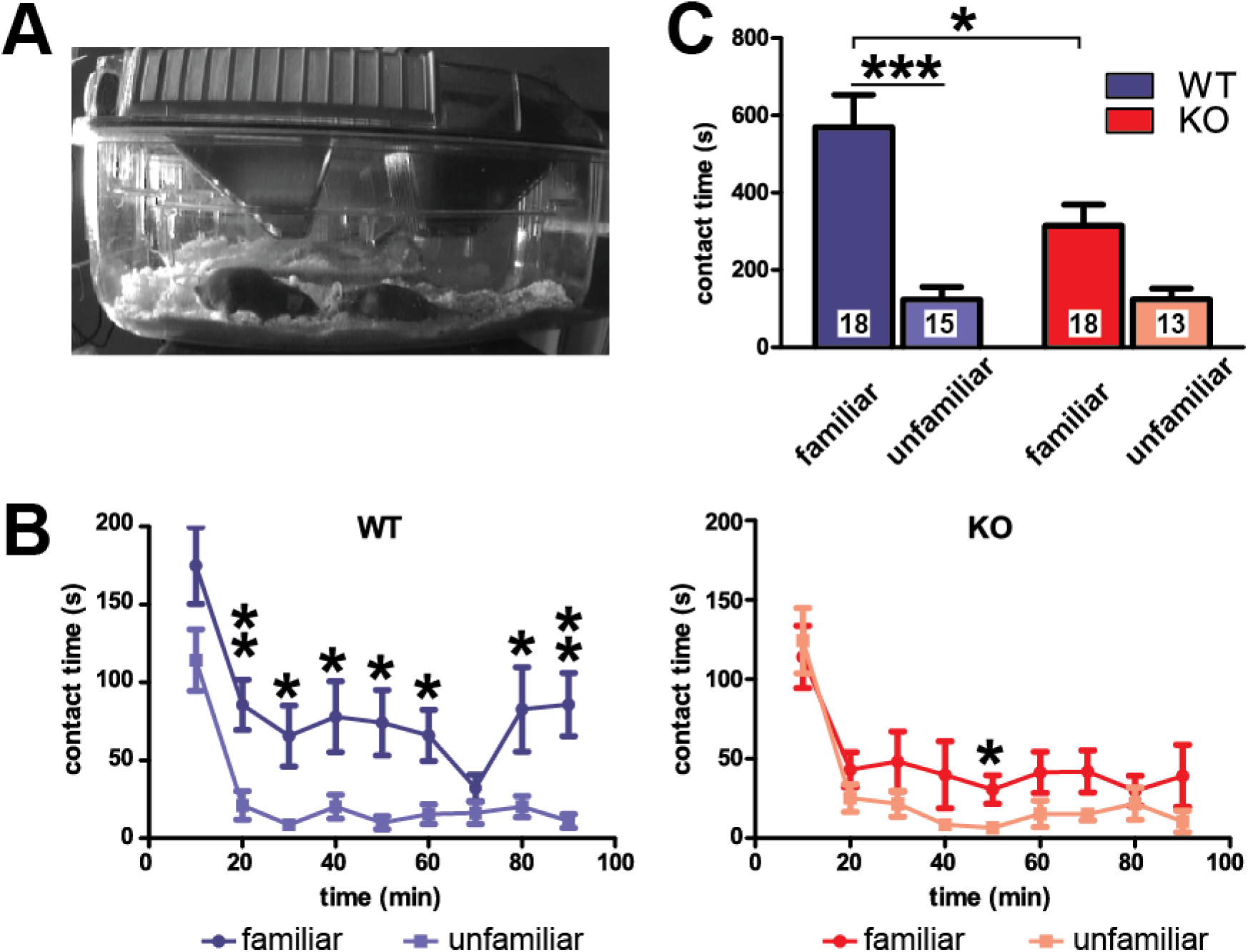
A decreased preference to interact with familiar anesthetized conspecific in BDNF KO mice. A) The interaction with anesthetized conspecific paradigm. A snapshot of a typical contact of a subject (left) with an anesthetized familiar conspecific (right). An infrared LED lamp illuminates the cage from the left side. B) Time courses for the duration of contacts made by WT (left, blue) and KO (right, red) subjects, shown in 10 min bins. The plots of darker and lighter colors correspond to the familiar and unfamiliar demonstrator, respectively. C) Summary bar diagram for the total durations of contacts with familiar and unfamiliar conspecifics by WT (blue) and KO (red) subjects. Numbers of animals are shown on the bars. Unpaired two-tailed t-test in B, Bonferroni post-hoc analyses in C: *p < 0.05, **p < 0.01, ***p < 0.001. Error bars represent s.e.m.

Total four independent groups of the KO and WT mice presented with either familiar or unfamiliar demonstrators were examined (Fig. 1). In all groups, robust contacts were observed during the first 10 minutes followed by the lower level but steady contacts during the remaining 80 min (Fig.1B). For the intense contacts during the first 10 min, there was no significant genotype*familiarity interaction or no significant main effect of either familiarity or genotype. During the subsequent 80 minutes, there was a significant genotype*familiarity interaction (F(1,60)=4.59, p=0.036) alongside a significant main effects of both familiarity (F(1,60)=28.6, p<0.001) and genotype (F(1,60)=4.58, p=0.038). In the time bin analysis along the 90 min observation period, WT mice exhibited a significantly longer duration of contacts with familiar mice during seven out of nine 10-min time bins, whereas KO mice showed a significant difference only in one bin (Fig. 1B). There were no differences between genotypes in contacting unfamiliar mice. For the entire 90 min, WT mice spent significantly more time contacting familiar stimuli (t=5.38, p<0.001), whereas KO mice only showed a tendency to do so (t=2.2, p>0.05) (Fig.1C). Together, these data indicate that when compared to the WT controls, the KO mice have a reduced preference to contacting familiar over unfamiliar anesthetized mice in the home cage.

### Effect of BDNF CA3 KO on non-social behaviors in the presence of familiar and unfamiliar demonstrator

Since the differences between WT and KO in contacting familiar demonstrators may reflect changes in non-social behaviors that compete with the contacting activity, we quantified the non-social behaviors at the time-resolution of a single video frame. The behaviors included eating from food hopper and drinking from water sipper (Eating), hanging from metal wire lid (Hanging), sitting still or sleeping alone in cotton nest (Resting in Nest), digging wood bedding (Digging), self-grooming (Grooming), rearing (Rearing), sitting still outside of the cotton nest (Not Moving).

First, before the detailed ethology, we analyzed locomotion of the subjects during the test. For the total distance traveled, the two-way ANOVA did not detect a significant genotype*familiarity interaction or a significant main effect of familiarity or genotype. For the average distance to the demonstrator, the ANOVA detected a significant main effect of familiarity (F(1,60)=8.06, p=0.006) and the Bonferroni posttest revealed a significantly longer average distance to the unfamiliar demonstrator in the WT group (t=2.48, p<0.05) (Fig.2A-C). There were no significant effects on the durations of Digging, Grooming, Rearing, and Not Moving (Fig.2D lower row panels) but in the presence of unfamiliar demonstrator, there were opposing tendencies in WT and KO mice towards more and less Digging, respectively, and a tendency towards more Not Moving in both genotypes.

**Fig. 2.**
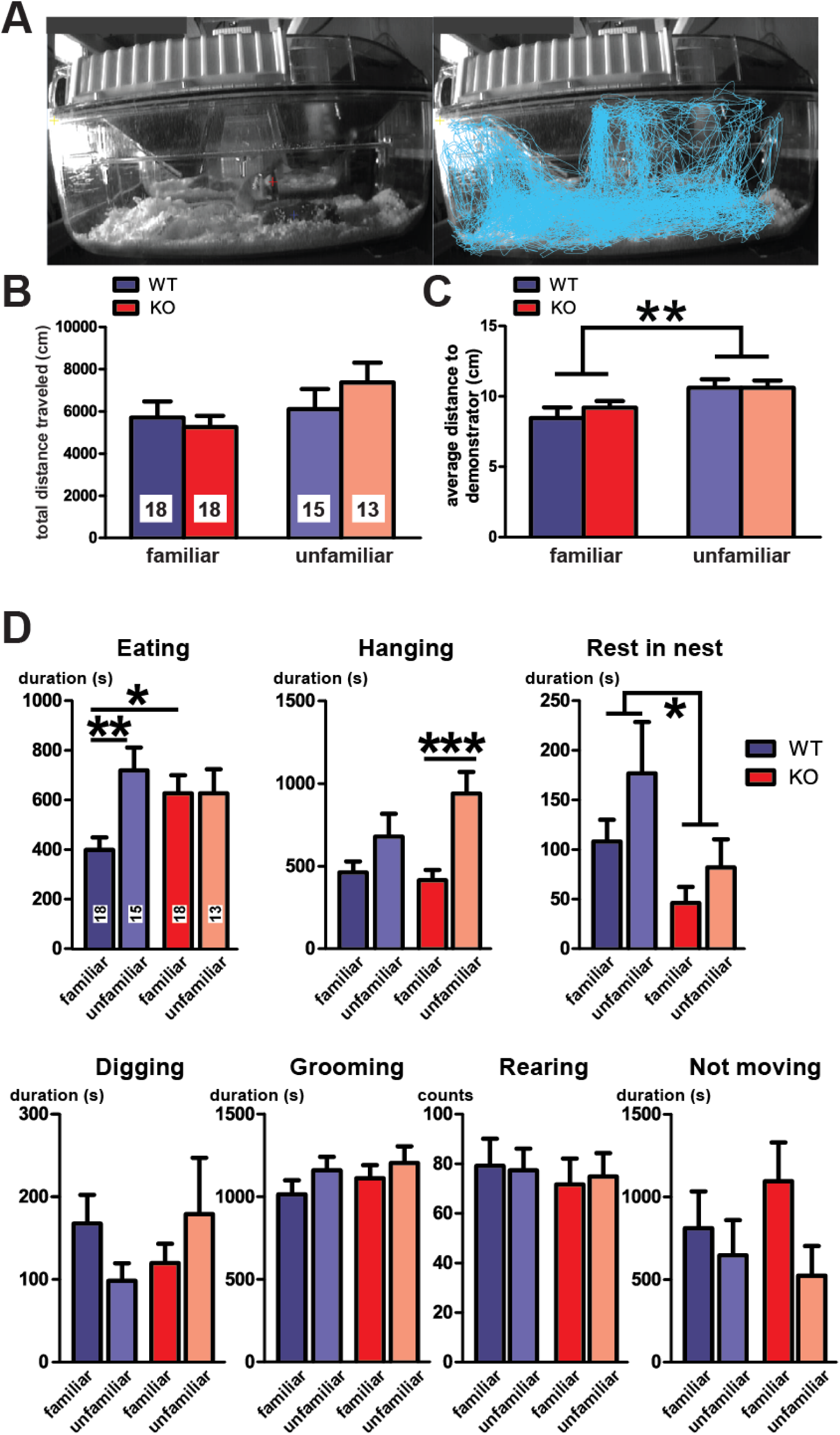
Non-social behaviors during the interaction with anesthetized conspecific. A-C) Genotype of subject and familiarity of demonstrator do not affect locomotion. A) Example pictures without (left) and with the overlay of moving trajectory (right) during the 90 min session with an anesthetized conspecific. B-C) Summary diagrams for total distance traveled and average distance to the demonstrator. D) Summary diagrams for the duration of other non-social behaviors. Blue and red colors represent WT and KO subjects, respectively. Cage mates (familiar, represented by darker colors) and 129 background mice (unfamiliar, represented by lighter colors) were used as the anesthetized demonstrators. Numbers of animals are shown on the bars. ANOVA, main effect of demonstrator in C and Bonferroni post-hoc analyses in D: *p < 0.05, **p < 0.01, ***p < 0.001. Error bars represent s.e.m.

By contrast, for Eating, there was a significant genotype*familiarity interaction (F(1,60)=4.2, p=0.044). The Bonferroni post-hoc analyses revealed that the WT mice spent significantly less time eating in the presence of familiar than unfamiliar demonstrator (t=2.97, p<0.01), whereas the KO mice did not (t=0.003, p>0.05). However, there was no significant negative correlation between Eating and contacting demonstrator (r=−0.043, p=0.73), which indicated that these two behaviors did not compete. For Hanging, there was no significant interaction between the two factors but a significant main effect of familiarity (F1,60)=14.2, p<0.001). Post-hoc analyses revealed that the KO mice spent significantly more time hanging in the presence of unfamiliar demonstrator (t=3.7, p<0.001), whereas the differences in WT mice were not significant (t=1.6, p>0.05). For Resting in Nest, there was a significant main effect of genotype (F(1,60)=6.3, p=0.015) but no genotype*familiarity interaction. Together, these data indicate that the genotype of subjects and the familiarity of anesthetized demonstrator influence several non-social behaviors without altering the overall activity of the subject.

### Normal sociability of BDNF CA3 KO mice

Sociability, or a propensity to spend time with another awake animal (Moy et al., 2004), was examined as a trait that could relate to the decreased contacting of KO mice with the anesthetized demonstrator. The three-chamber sociability task (Moy et al., 2004) was conducted on two groups per each genotype using either familiar (cage mates) or unfamiliar (age-matched 129SvEv background male mice) awake demonstrator in the cup. With either type of demonstrator, the subjects spent more time near the cup containing demonstrator vs an empty cup (unfamiliar: F(1,33)=7.8, p=0.009; familiar: F(1,26)=24.6, p<0.0001) but there was no significant cup*genotype interaction (unfamiliar: F(1,33)=0.25; p=0.62; familiar: F(1,26)=0.005, p=0.94) (Fig.3). In both genotypes, the post-hoc analyses revealed significant preference towards spending more time with familiar demonstrator than with empty cup (WT: t=3.7, p<0.01; KO: t=3.3, p<0.01), whereas, with unfamiliar demonstrator, the preference did not reach significance (WT: t=1.7, p=0.07; KO: t=2.2, p=0.06). In addition, the two-way ANOVA did not detect a significant stimulus (familiar vs unfamiliar)*cup (containing demonstrator vs empty) interaction. Together, these data indicate that KO and WT mice have the same level of sociability.

**Fig. 3.**
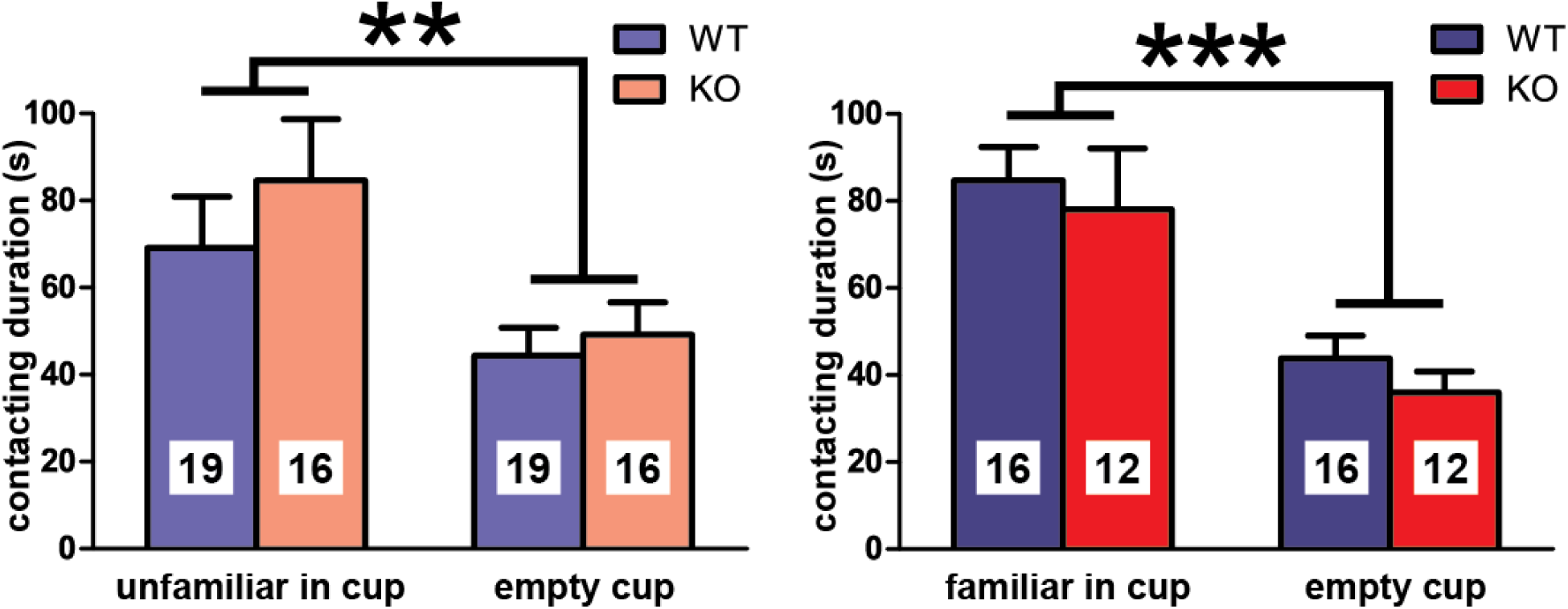
Normal sociability in KO mice. Summary diagrams for contact durations with empty cup or cup with a demonstrator, either unfamiliar (left) or familiar (right). Blue and red colors represent WT and KO subjects and the number in each bar indicates the number of subjects. ANOVA, main effect of cup: **p < 0.01, ***p < 0.001. Error bars represent s.e.m.

### Olfaction of social and non-social odors in KO mice is normal

Olfaction is the major sensory modality that drives social behaviors in rodents (Arakawa et al., 2008). Since BDNF KO male mice have normal social recognition (Ito et al., 2011), it is less likely that an impaired recognition of familiarity prevented KO mice from changing the contacting and eating activities. Nevertheless, we tested for a potential olfactory deficit in KO mice using the olfactory habituation/dishabituation test (Crawley et al., 2007) with two non-social and two social odors (Fig.4). Odor habituation was defined as a decline of sniffing duration along three consecutive presentations of an identical odor. It was significant in both genotypes with all odors (WT, F(2,19)>6.0, p<0.005; KO, F(2,22)>8.9, p<0.001) except for the WT mice with the odors of lemon (F(2,19)=3.0, p=0.064) and urine from 129SvEv males (F(2,19)=0.5, p=0.61). Odor dishabituation was defined as an increase in sniffing duration upon presentation of a new odor. The dishabituation was significant in both genotypes with all odors (WT, F(2,19)>8.6, p<0.009; KO, F(2,22)>9.3, p<0.006), except for the WT mice presented with the lemon odor (F(2,19)=4.2, p=0.056). There was no odor*genotype interaction in the habituation/dishabituation tests. These findings indicate that KO mice have normal olfaction.

**Fig. 4.**
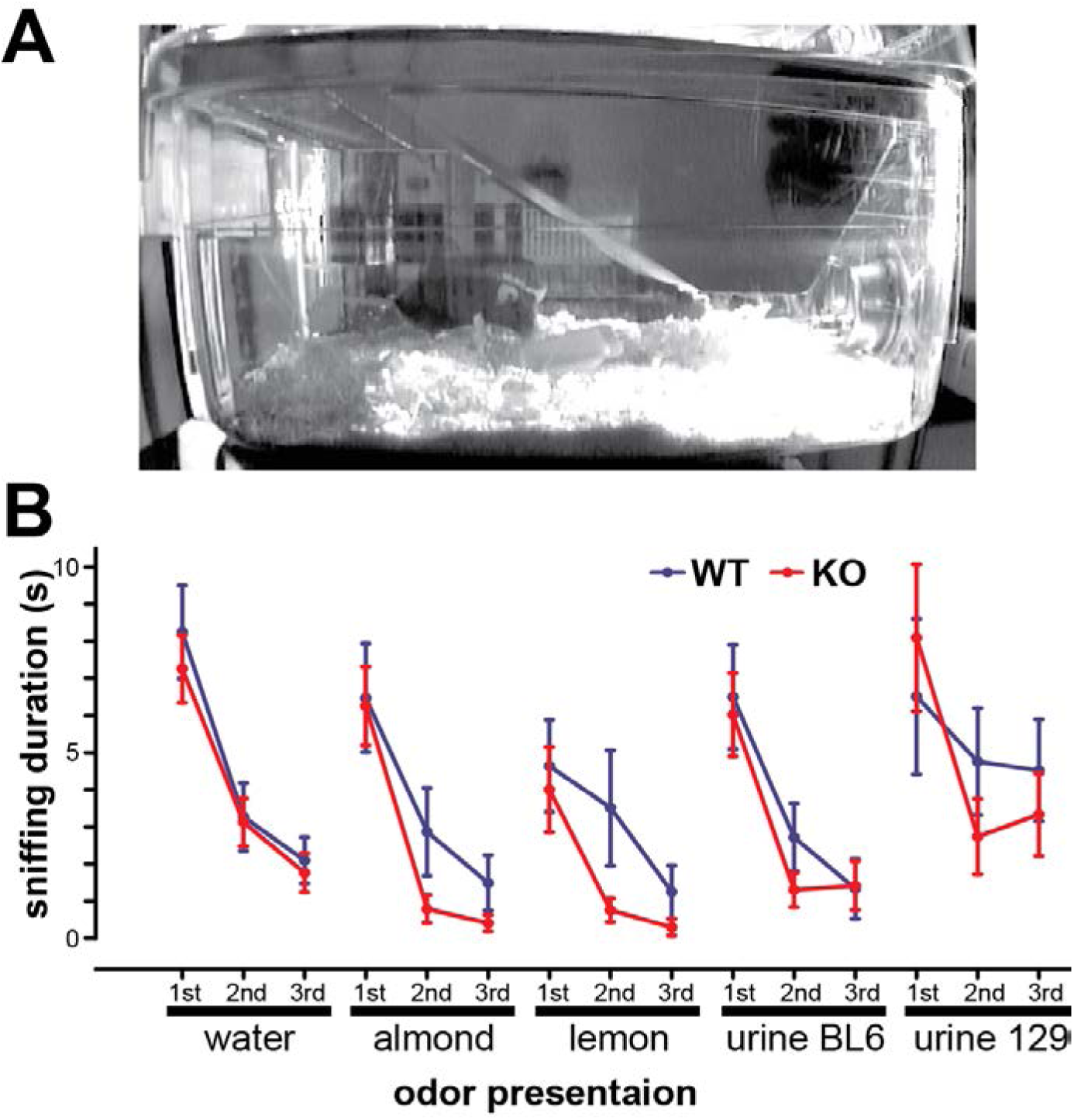
Olfactory habituation/dishabituation test. A) An example of an animal sniffing an odor presentation tray with a novel odor. B) Summary diagram for the sniffing duration in WT (blue) and KO (red) mice. The X-axis shows the sequence of odors’ presentations. n= 20 WT and 23 KO mice. Error bars represent s.e.m.

### Blunted pain sensitization in BDNF KO mice

Although BDNF KOs distinguish odors including urine smells from conspecifics of different genetic backgrounds, the failures to sustain contacts with anesthetized cage mate and to decrease eating may result from an inability to perceive the state of others or a deficit in empathy-like behaviors. Given the overlap between neuronal pathways implicated in such perception and the pathways involved in pain sensitization (Engen and Singer, 2013; Li et al., 2014), the responses to persistent pain were examined using the formalin test.

Upon formalin injection, WT mice exhibited typical biphasic nociceptive response (Bannon and Malmberg, 2007) (Fig.5), with intense paw licking during the first five minutes after formalin injection, followed by a decline and then by the second phase with the licking peak at 15-25 min after the injection. The KO mice exhibited same levels of licking with the WT mice during the first five minutes (phase 1) but significantly less licking during the 15-25 min time interval (phase 2) (p<0.01, t-test), which suggests that KO mice have normal response to acute pain but are impaired in sensitization to the persistent pain.

**Fig. 5.**
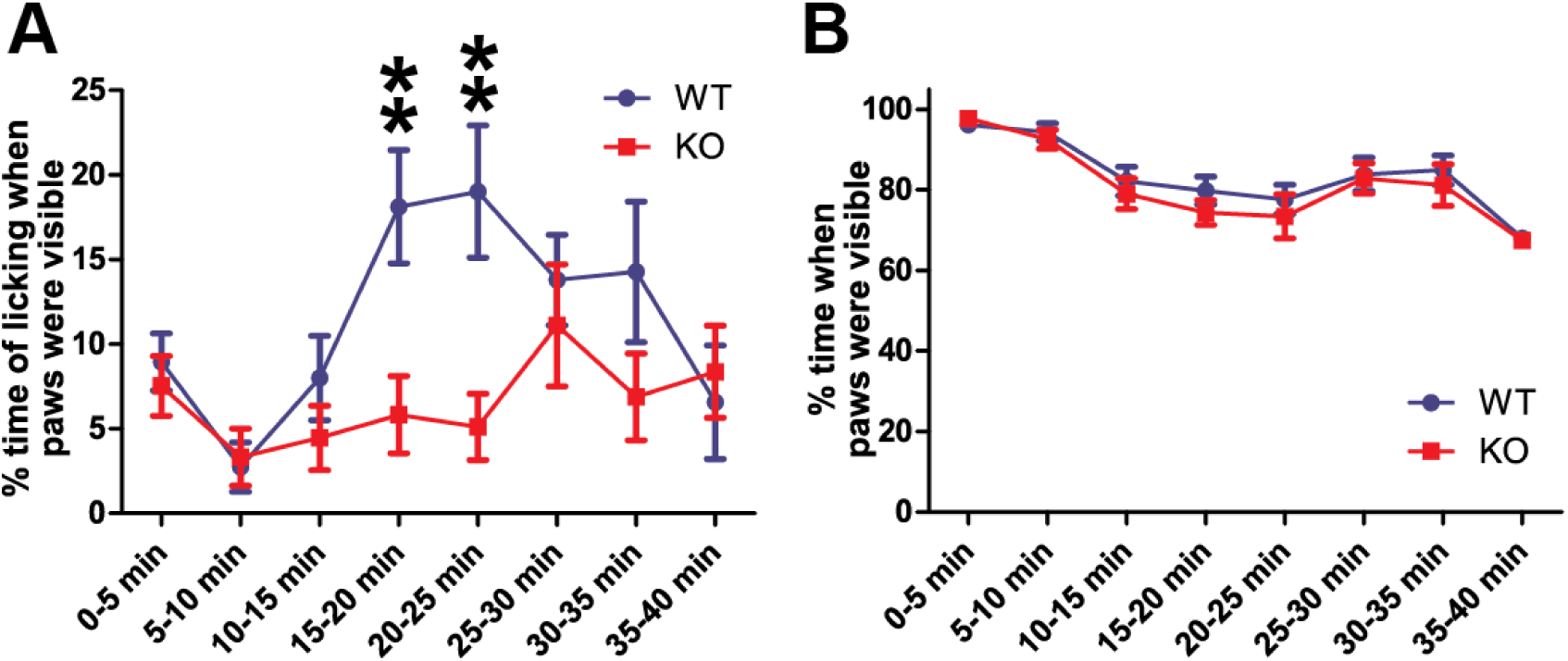
Blunted pain sensitization in KO mice. A) Summary diagram for paw licking time expressed as the percentage of the total time when animal paws were clearly visible during 5 min bins along the 40 min after formalin injection. B) Percentages of time when animals’ paws were clearly visible. Blue and red represent WT (n=18) and KO (n=15) subjects. Unpaired t-test: **p < 0.01. Error bars represent s.e.m.

## Discussion

Here, we report a novel social trait in mice - the preference towards making repeated social contacts with a familiar over unfamiliar anesthetized conspecific in the home cage environment. Then, we find this trait compromised in mice with the CA3-restricted knockout of BDNF, whose high aggression against an awake conspecific (Ito et al., 2011) masks more subtle social behaviors.

There are two advantages of using anesthetized demonstrator for investigating social behaviors. First, the subject activity is not affected by the behavior of demonstrator. In contrast, with awakened demonstrator, social contacts are initiated and terminated not only by subject but also by a demonstrator, which makes it difficult to attribute the pattern and amount of social interaction solely to the properties of the subject. This problem is partially solved by restricting movement of the demonstrator to a wire cup (Moy et al., 2004), which makes it possible to isolate active social touches made by the subjects but does not entirely exclude influences from the demonstrator. Second, the anesthetized demonstrator is less likely to induce aggression, which requires a chain of reciprocal activities (Ito et al., 2011) and potentially masks other social behaviors. Conversely, the limitation of the test is the omission of social behaviors driven by reciprocal interactions.

In this study, all subjects did not express aggression but actively contacted demonstrator regardless whether it was familiar or not. During the first ten minutes of the test, when there was a burst of contacting activity, the familiarity of demonstrator did not affect the duration of contacts. However, during the following 80 minutes, when the overall level of contacts decreased, animals spent more time contacting the familiar demonstrator. It suggests that while the initial highly intense contacts are driven by exploration and novelty seeking, the subsequent contacts are driven more by the social cues that are already familiar to the subject and that those familiar cues enable the sustained contacting activity (Fig. 1). The sustained contacts with familiar animals did not involve huddling. However, when demonstrator woke up and began to move, the huddling returned, which indicates that huddling requires social signals from an awakened animal. The lack of a significant effect of genotype on the initial contacts suggests that the exploratory drive and novelty seeking are not compromised by the CA3 BDNF deletion. The normal response to novelty was also supported by the normal olfactory dishabituation upon presentation of novel odors (Fig.4). By contrast, the significantly decreased duration of the later contacts suggests a deficit in processing the familiar social cues.

How the familiar social cues cause sustained and relatively high contacting activity? Novelty seeking, sexual drive or aggression against a competing animal does not explain the contacts. One explanation could be the drive to affiliate with a familiar conspecific (Panksepp et al., 2007), but it appears contradicting the natural preference of mice for social novelty (Moy et al., 2004). An alternative but intriguing idea is that the irregular state of the anesthetized familiar conspecific is the cause. The anesthetized cage mate generates social cues recognized as familiar by the subject but does not express predicted behaviors, even upon social contacts. The conflict between predicted and observed behaviors could trigger the elevated contacting activity and possibly suppress feeding. Ethologically relevant situations could be the encounters with a sick, injured or distressed conspecific. The stronger response to the lack of expected behaviors from a familiar versus an unfamiliar animal may indicate a higher sensitivity to the state of the partner than of a stranger and be, therefore, categorized as one of the empathy-like traits, for some of which the familiarity is the major determinant (Jeon et al., 2010; Langford et al., 2006).

Then, what is wrong in BDNF KO mice? Their decreased preference to contacting familiar demonstrator could not be explained by changes in sociability, which was found normal. Neither it could be explained by a failure to recognize a cage mate because these mice have normal social recognition (Ito et al., 2011) and the ability to recognize social and non-social odors (Fig.4). A possible blunted sensitivity to the state of the partner, however, could explain not only the reduced contacting time with the anesthetized familiar demonstrator but also the inability to reduce aggression against a cage mate even when it shows submission (Ito et al., 2011). In addition, the atypical response of KO mice in the formalin test suggests a malfunction of the neuronal mechanisms underlying sensitization to persistent pain, possibly in the anterior cingulate cortex (Zhuo, 2007), which has been implicated in empathy in humans and empathy-like behaviors in rodents (Engen and Singer, 2013; Jeon et al., 2010; Li et al., 2014). While the link between the hippocampal CA3-restricted BDNF knockout and neuronal mechanisms of empathy-like behaviors has not been established, the evidence that normal late development of the prefrontal cortex requires the hippocampus (Bertolino et al., 2002; O’Donnell et al., 2002), points to a possibility that the loss of hippocampal BDNF is causing a relevant malfunction in the prefrontal cortex.

## Materials and Methods

### Animals

Mice with the CA3-restricted knockout of BDNF were generated by combining two mutant mouse lines, the floxed BDNF line (Zakharenko et al., 2003) and the transgenic bacterial artificial chromosome KA1 Cre recombinase driver line (Nakazawa et al., 2002) as previously described (Ito et al., 2011). Prior to interline crossings, these lines were backcrossed to C57BL/6 background animals a minimum of 6 generations. To produce animals for experiments, homozygous BDNF-floxed Cre-positive (BDNF^ff, Cre^) males were crossed with homozygous BDNF-floxed Cre-negative (BDNF^ff^) females to obtain BDNF^ff, Cre^ and BDNF^ff^ male animals, further referred to as knockout (KO) and wild type (WT), respectively. The genotype of mice was determined as previously described (Zakharenko et al., 2003).

### Behavior

All experiments were approved by Virginia Tech IACUC and followed the NIH Guide for the Care and Use of Laboratory Animals. Male mice were weaned around p21-p25 when body weight exceeded 10 g and housed as pairs of littermates of the same genotype in a regular 12:12 h dark-light cycle. Bedding was hardwood chips (Beta Chip, NEPCO, Warrensburg, NY). A TP roll (1.5"×4.5", Jonesville Paper Tube Corp, Jonesville, MI), a 2"×2" Nestlet, and a pinch of Enviro-Dri (PharmaServ, Framingham, MA) served as environmental enrichment. Food (Rodent NIH-07 open formula diet, Envigo, Cambridgeshire, UK) and water were provided ad libitum. Experiments were performed at p40–p60, prior to the onset of aggression toward cage mate (Ito et al., 2011). Behavioral experiments were done during the light phase of the light-dark cycle under the illumination of 200 lux except for the interaction with anesthetized conspecific. The days of weekly cage changes were avoided.

#### Interaction with anesthetized conspecific

The experiments were performed using the home cages that housed subject mice for no less than two days after cage change. A cage mate or an age-matched 129SvEv male mouse (a demonstrator mouse) was anesthetized with intramuscular injection of ketamine/xylazine/acepromazine (100/20/3 mg/kg) and placed at the center of the cage. When a 129 mouse served as a demonstrator, the cage mate was removed prior to introducing demonstrator. The behavior of the subject was recorded digitally at 5-8 frames per second (fps) using the StreamPix5 software (Quebec, Canada) in a dark room under infrared LED illumination. The sessions started at the beginning of the dark cycle (7 pm) and lasted for 90 min. Beginning and end of each epoch of body contact (defined in the results) between subject and demonstrators were determined offline by experimenters blind to the animal genotype using a custom-made behavior annotation module for the StreamPix5 software, which allows annotations for predefined behaviors at the resolution of a single video frame. In addition, the durations of eating, drinking, hanging from the wire lid, sitting still inside the cotton nest, digging bedding, self-grooming, not moving outside the nest and the counts of rearing were determined using the same method. To quantify locomotion inside the home cages, the trajectories of animal movements were tracked manually using a custom-made tracking module for StreamPix5 software and a pen tablet connected to a PC.

#### The sociability test

The sociability test was performed as described (Moy et al., 2004). The sociability chamber (60 × 40 cm, ANY-maze, Wood Dale, IL) made with transparent Plexiglas sheet consisted of three compartments (20 × 40 cm) connected by two gates (width × height: 5 × 8 cm) with sliding doors. Two small, round wire cups (black, diameter × height: 10.5 × 11 cm, Galaxy Pencil & Utility Cup, Spectrum, Streetsboro, OH) were placed at the centers of both side compartments. Plastic cups (SOLO^®^ Plastic Cold Party Cups, Red, 16 oz) filled with water were placed on the top of the wire cups to prevent the subject from climbing the wire cups. The behavior of the subject was recorded using the Streampix5 software and 3 digital video cameras, viewing from the top and from each side of the chamber to avoid any blind spots. The experimenter hid behind a curtain. The subject was first acclimated in the center compartment with doors closed for 5 min. Then, a demonstrator mouse was introduced into one of the cups selected randomly and the doors were opened. The subject was allowed to explore the compartments for 10 min. Cage mates and age-matched 129SvEv mice were used as familiar and unfamiliar demonstrators, respectively, and were acclimated within 1-2 days before the test by being placed inside the wire cup for 30 min. The beginning and end of the behavior epochs when subjects were attending towards the cups or were physically touching them were annotated the same way as the interaction with anesthetized conspecifics.

#### Olfaction test

The olfactory habituation/dishabituation test was performed as described (Yang and Crawley, 2009) with slight modifications. Weighing boats (4.5 × 4.5 cm) with a piece of Whatman paper (2 × 2 cm) attached by Scotch double sided adhesive tape were used for odor presentation. On day 1, subject animals were acclimated for 1 hour to the test environment, which was a clean cage with a metal lid, a cover top and a clean odor presentation boat placed on fresh wood bedding. The cage was located on a rack equipped with monitoring cameras. On day 2, after 30 min acclimation, sequential presentation of odors was performed repeatedly 3 times for each odor, using freshly prepared odor presentation boats spotted with 10 μL of water, imitation almond (1:20 dilution, Kroger), lemon extract (1:20 dilution, Por Han-Dee Pak, Inc), urine from C57BL6 males and urine from 129SvEv males. One presentation consisted of 2 min placement of the boat at the center of the cage. The interval between presentations was around 1 min. The animal behaviors were recorded using digital video cameras and StreamPix5, while the experimenter hid behind a curtain. Although the room illumination was set at 200 lux, an IR LED illumination was applied from the side of cages for optimal video recording. The analysis was performed offline by persons unaware of the subject genotype. The beginning and the end of the subject sniffing the boat were recorded. The subjects that buried the boat by digging around it were excluded from the analysis.

#### Formalin test

The test was performed as described (Bannon and Malmberg, 2007; Dubuisson and Dennis, 1977) in transparent plastic cylinders (diameter × height: 13 × 15 cm) positioned within compartments to prevent animals in neighboring cylinders from seeing one another while allowing videotaping from the front. Two mirrors were assembled at the angle of 90 degrees and placed at the back of each cylinder to maximize the visibility of the subject. The experimenter hid behind a curtain. On day 1, each subject was acclimated to the cylinder for 30 min. On day 2, mice received subcutaneous 20 μL injections of 5% formalin in the middle of the hind paw on the plantar side using a 50μ Hamilton syringe with a 30G needle. The animals were placed in the cylinder and videotaped for 40 min using the StreamPix5 software. The recordings were analyzed offline using the annotation module for StreamPix5 by experimenters blind to the animal genotype. The duration of epochs when animal licked the hind paw and when the hind paws were invisible by the camera was determined.

### Statistics and data analysis

Two-way ANOVA and the Bonferroni post-hoc test were used to analyze interaction with the anesthetized demonstrator, non-social behaviors, and sociability in the three-chamber experiment. The factors were genotype, demonstrator (familiar or not) and cup (empty vs with demonstrator). The Student t-test was used to compare contact times with familiar and unfamiliar anesthetized demonstrator during the 10 min time bins and to compare licking times in the formalin test. Two-way repeated measure ANOVA was used in the olfactory habituation/dishabituation test with odor and genotype as the factors. The correlation between contacting and eating was tested using the Pearson correlation coefficient. Effects were considered significant at p<0.05.

## Acknowledgements

The study was supported by NIH grant K22MH097826.

## Competing interests

Authors declare no competing interests.

